# Identification of miR136, miR155, and miR183 in vascular calcification in human peripheral arteries

**DOI:** 10.1101/2024.10.09.616039

**Authors:** Tom Le Corvec, Mathilde Burgaud, Marja Steenman, Robel A Tesfaye, Yann Goueffic, Blandine Maurel-Desanlis, Thibaut Quillard

**Affiliations:** CHU Nantes, L’institut du thorax, service de chirurgie vasculaire, Nantes, France; CHU de Clermont-Ferrand, Hôpital Gabriel Montpied, Clermont-Ferrand, France; Nantes Université, CHU Nantes, CNRS, INSERM UMR 1087, l’institut du thorax, F-44000 Nantes, France; Laboratory of Chromatin Dynamics, Centre de Biologie Intégrative (CBI), MCD Unit (UMR5077), Université de Toulouse, CNRS, UPS, 31062 Toulouse, France; Groupe Hospitalier Paris St. Joseph, Department of Vascular and Endovascular Surgery, Paris, France

**Keywords:** Atherosclerosis, Vascular Calcification, miRNAs, Vascular Smooth Muscle Cells

## Abstract

**Objective:** Vascular calcification (VC) is an independent risk factor for all-cause and cardiovascular mortality. This process contributes to atherosclerotic plaque disruption and thrombosis when close to the lumen, arterial stiffness, and limits endovascular treatment success. Vascular smooth muscle cells (VSMC) in the arterial wall play a major role in VC as they can acquire mineralizing properties when exposed to osteogenic conditions. Despite its clinical impact, there are still no dedicated therapeutic strategies targeting VC.

**Design:** To address this question, we used human calcified and none-calcified atherosclerotic arteries (ECLAGEN Biocollection) to screen and identify microRNA (miRs) associated with vascular calcification.

**Methods:** We combined non-biased miRNomic (microfluidic arrays) and transcriptomic analysis to select miR candidates and their putative target genes, with expression associated with vascular calcification and ossification. We further validated miR functional regulation and function on cell mineralization using primary human vascular SMC.

**Results:** Our study identified 12 miRs associated with vascular calcification in carotid and femoral arteries. Among those, we showed that miR136, miR155, and miR183 expression were regulated during VSMC mineralization and overexpression of these miRs was sufficient to promote smooth muscle cell mineralization. Cross-analysis of this miRNomic and a transcriptomic analysis led to the identification of CD73 and Smad3 pathways as putative target genes responsible for mediating miR155 pro-mineralizing function.

**Conclusion:** These results highlight the potential benefit of miR155 inhibition in limiting VC development in peripheral atherosclerotic arteries.

**What this paper adds:** Vascular calcification (VC) is an independent risk factor for all-cause and cardiovascular mortality. No pharmacological treatment is currently available. Our work using human healthy and atherosclerotic peripheral arteries allowed us to non-biasedly identify miR136, miR155, and miR183 as putative players in VC, as they associate with calcification in human atherosclerotic lesions, and are sufficient to induce vascular cell mineralization *in vitro*. Among those, miR155 appeared the most important driver of osteogenesis *in vitro*, making it a candidate for targeting VC in patients.

## Introduction

Vascular calcification (VC) of peripheral arteries is an independent risk factor for all-cause and cardiovascular morbi-mortality^1–3^. The risk of mortality can also differ depending on vascular beds affected by VC^4^. Intimal Calcification is the main form of VC and consists of calcification development within atherosclerotic lesions. The composition and nature of calcification can affect plaque stability. While microcalcifications in proximity to the lumen are linked to an elevated risk of plaque rupture and acute events, recent research indicates that in the context of coronary arteries, intimal macrocalcifications may serve to locally stabilize vulnerable lesions^5,6^. Furthermore, arterial stiffness resulting from VC represents an additional risk factor for morbi-mortality, as it contributes to the development of hypertension and heart failure. Additionally, the protrusion of calcified nodules within the lumen can directly facilitate thrombosis^7^. Ultimately, VC has a major direct negative impact on clinical practice, by jeopardizing surgical or endovascular treatments and directly affecting the success of interventions and prognosis in patients with peripheral artery diseases (PAD)^8–10^.

The development of vascular calcification is the result of an imbalance in pro-calcifying factors in pathological conditions such as atherosclerosis and chronic kidney disease. Different mechanisms leading to VC, including high phosphorus concentration, cell death, and inflammation, can prompt VSMCs to transdifferentiate into osteoblastic properties and overexpress genes like Runx2, Sox9, Osteopontin, Osteocalcin, and TNAP^11,12^. Conversely, VC inhibitors like pyrophosphate (PPi) are decreased under these pathological conditions, through a lower production by ENNP-1 and/or increased conversion into inorganic phosphate by alkaline phosphatase TNAP^13^.

Previous studies conducted by our research team using human arteries (ECLA and ECLAGEN biocollections) have demonstrated that arteries develop heterogeneous lesions and mineralization between vascular beds^14^. Arterial plaques in femoral arteries exhibit a higher calcium content and are more prone to bone-like calcification. In vitro studies have shown that femoral vascular smooth muscle cells (VSMC) have a higher propensity for mineralization than other territories^11^. The molecular specificities among vascular beds, especially in non-atherosclerotic arteries, imply that vascular calcification types could be driven by inherent molecular variability between vascular cells from these locations^15^. In light of these considerations, we sought to identify microRNAs (miRs) associated with vascular calcification in the carotid and femoral arteries. We employed a similar approach with our transcriptomics data to identify the target genes through which miRs might regulate these processes.

The post-transcriptional regulation of gene expression by miRs has demonstrated its significance in human diseases and vascular calcification of coronary arteries^16,17^. Some miRs have a protective effect in inhibiting VSMCs osteoblastic trans-differentiation (miRs34- b, 205 and 26-a)^18–20^ and by regulating Beta Glycerophosphate/Runx2 pathway (miR204 and 140-5p)^21,22^. Conversely, overexpression of miRs17-5p, 29 and 221/222) affects regulatory pathways that promote VC^23–25^.

Identification of miRs regulating VC of peripheral arteries is important to improve our knowledge and further explore new therapeutic strategies. The aim of this work was an unbiased identification of miRs associated with VC in human carotid and femoral arteries, to characterize the involvement of such miRs in VSMC mineralization, and to determine their target genes and functional role.

## Material and Method

Comprehensive material and methods (RNA extraction, miRs microfluidic arrays, and transcriptomic analysis; Cell culture and mineralization; Cell transfection and functional validation of interest miRs; Gene and miR expression analysis; In silico analysis of the miR candidates; Protein expression analysis) are described in supplementary material, provided online.

### Biological samples

ECLAGEN biocollection includes human carotid bifurcation and common femoral artery samples. Details of these biocollections have been described in previous publications^26,27^. Pathological atheromatous tissues were collected during endarterectomy and healthy arteries were collected from organ donors. In the operating room, the atheromatous lesions coming from the endarterectomy were cut into two pieces, one for RNA extraction and molecular analysis, and the other for histological analysis. Stratification of lesions calcification was performed after histomorphometry analysis of mineralized structures as depicted in our previous studies^11^. The collection of human samples was carried out according to strict ethical standards. All living participants received an information notice and signed a written consent (research protocol#PFS09-014, authorized on December 23, 2009, by the French “Agence de Biomédecine”). For deceased donors, no opposition to organ donation was checked, and written consent from the donor’s family was obtained, therapeutic arterial samples were always prioritized. The legal authorizations were obtained from the French Ministry of Research (n DC-2008-402), the National Commission for Informatics and Freedoms (CNIL, n 1520735 v 0), and the Local Ethics Committee (GNEDS, Groupe Nantais d’Ethique dans le Domaine de la Santé). Histological study of this biocollection had been carried out in previous studies^11,15^.

### RNA extraction, miRs microfluidic arrays, and transcriptomic analysis

Expression of 753 human miRs was carried out with micro-fluidic cards (TaqMan™ Array Human MicroRNA cards, ThermoFisher). The analysis was conducted on 60 arterial samples from ECLAGEN biocollection: 20 atherosclerotic carotid arteries (10 calcified, 10 none calcified), 20 atherosclerotic common femoral arteries (10 ossified, 10 none ossified), 10 healthy carotid arteries and 10 healthy common femoral arteries. Adventitia was removed before RNA extraction. Total RNA extraction from tissue and primary cells was performed using Rneasy Plus Micro kit by following the manufacturer’s instructions (Qiagen). For tissue samples, Tissuelyzer II (Qiagen) was used for homogenization before RNA extraction.

Following the completion of the Qiazol and column-based extraction procedure, RNA concentration was determined by spectrophotometry (NanoDrop 1000, ThermoFisher). The Reverse Transcription was achieved with the RT Maxima H Minus Enzyme Mix kit according to the manufacturer’s recommendations (ThermoFischer). mRNA analysis was performed on microarray data from the same samples as previously reported^21^,deposited in NCBI’s Gene Expression Omnibus (GSE100927). Gene expression differences between calcified and non-calcified atherosclerotic carotid arteries, and between ossified and non-ossified femoral arteries were analyzed using the limma R/Bioconductor software package (Ritchie et al. NAR 433(7), 2015). The obtained results were visualized using the VolcaNoseR web app (Goedhart & Luijsterburg, Scientific Reports 10:20560, 2020).

### Cell culture and mineralization

Primary’s Human Aortic Smooth Muscle Cells were isolated from human aorta (PromoCell, 3 separate donors) and cultured in a dedicated proliferation medium (Smooth Muscle Cell Growth Medium 2, PromoCell), including fetal calf serum (5%), growth factors (Epidermal Growth Factor, Basic Fibroblastic Growth Factor) and insulin (5 µg/ml). Penicillin and Streptomycin (PS) were added to the 1% concentration (Sigma Aldrich).

VSMCs were cultured in two pro-mineralizing αMEM medium (Dulbecco’s Modified Eagle’s Medium) containing both high glucose (Gibco, Thermofisher). We added fetal calf serum (3%) and inorganic Phosphate at the concentration of 3 mM (“Pi-enriched Medium”) by adding NaH_2_PO_4_ (Molecular Sigma Biology) and Na_2_HPO_4_ (Fluka BioChemika) solutions (4:1 ratio). For the second medium (“OB medium”) we added fetal calf serum (10%), Dexamethasone 10^-7^ M and Vitamin D3 10^-8^ M (from day 4 to day 21), and β- Glycerophosphate 10 mM and Ascorbic acid 0.25 mM (from day 8 to day 21). Cells were then fixed with absolute alcohol for 20 minutes at 4°C. Mineralization staining was performed with Rouge Alizarine (Sigma Aldrich) for 20 minutes at room temperature. Stained wells were individually photographed with a binocular magnifying glass (Zeiss Stemi 200-C and AxioVision Rel 4.8 software). Mineralization was quantified with Image Pro Plus software by calculating the ratio of the mineralized area (stained in red) to the total well area.

### Cell transfection and functional validation of interest miRs

miR overexpression with mimic miRs (30nM, Qiagen) was achieved by lipofection (Lipofectamine RNAiMax, ThermoFischer). For mineralization experiments, the pro- mineralizing medium was added 24 hours after transfection and repeated every 7 days.

### Gene and miR expression analysis

Transcriptional analysis was obtained by Real-Time qPCR on the CFX96 (Bio-Rad) detection device with Power Sybr green PCR Master Mix (ThermoFisher). Target gene expression was normalized to GAPDH expression, and the comparative cycle threshold (Ct) method was used to calculate the relative expression of target mRNAs. Primers of target mRNAs were : *Glyceraldehyde-3-Phosphate Dehydrogenase (GAPDH)* R–GGTGCAGGAGGCATTGCT/F– TGGGTGTGAACCATGAGAAGTATG, *Runt Related Transcription Factor 2 (RUNX2)* R– GCTCTTCTTACTGAGAGTGGAAGG/F–GCCTAGGCGCATTTCAGA, *Osteocalcin (OCN)* R–GTGGTCAGCCAACTCGTCA/F–GGCGCTACCTGTATCAATGG, *Osterix (OSX)* R– GCCTTGCCATACACCTTGC/F–CTCCTGCGACTGCCCTAAT, *Osteopontine (OPN)* R– CAATTCTCATGGTAGTGAGTTTTCC/F–GAGGGCTTGGTTGTCAGC, *Collagen Alpha-1(I) Chain (COL1A1)* R–GCTCCAGCCTCTCCATCTTT/F–CTGGACCTAAAGGTGCTGCT,

*Alkaline Phosphatase (ALP)* R–GGTCACAATGCCCACAGATT/F– AACACCACCCAGGGGAAC, *Bone Sialo Protein (BSP)* R– CAGTCTTCATTTTGGTGATTGC/F–CAATCTGTGCCACTCACTGC, *Aggrecan (ACAN)* R– GACACACGGCTCCACTTGAT/F–CCCCTGCTATTTCATCGACCC, *SRY-Box Transcription Factor 9 (SOX9)* R–TCGCTCTCGTTCAGAAGTCTC/F-GTACCCGCACTTGCACAAC, *Actin Alpha 2 (ACTA2)* R–CCGGCTTCATCGTATTCCTGTT/F–TCCTTCATCGGGATGGAGTCT, *Tropomyosin 1 (TPM1)* R–CTCCTCTGCACGTTCCAGGT/F– AGGAGCGTCTGGCAACAGCT, *Transgelin (SM22-Alpha)* R–CACCAGCTTGCTCAGAATCA/F–CAGTGTGGCCCTGATGTG, *Calponin (CNN1)* R– GTACTTCACTCCCACGTTCACCTT/F–GAACATCGGCAACTTCATCAAGGC and *CD73* (QuantiTect N° QT00027279, Qiagen).

We analyzed miR expression by RT-qPCR using miRCURY LNA RT kit and miRCURY qPCR LNA kit (Qiagen). The miR primers were provided by Qiagen (LNA miRNA PCR Assay, Qiagen). The expression of miRs was normalized to miR SNORD 44 and 48 expression.

### *In silico* analysis of the miR candidates

We have determined the associated genes of the miR candidates using the miRWalk 2.0, miRbase, and miRBD databases^33^, and the main biological functions associated using the REVIGO online database (http://revigo.irb.hr/) and the gene ontology resource (http://geneontology.org/).

We performed a cross-analysis of the data from the transcriptomic analysis of ECLAGEN biocollection already published^16,21^ with the selected miRs and the gene ontology resource (http://geneontology.org/). We selected target genes of miR of interest found associated with calcified plaques and involved in the process of vascular calcification/ossification and/or cell mineralization.

### Protein expression analysis

Protein extraction was performed with RIPA lysis buffer containing protease inhibitors (P8340, Sigma Aldrich®). Quantification was conducted by spectrophotometric analysis using the BCA Protein Assay kit (ThermoFisher).

Protein expression analysis was performed by Western Blot after migration on Acrylamide 4 – 15% gradient gel with Bis-Tris buffer (Bio-Rad®). The transfer was then realized on a nitrocellulose membrane TransBlot Turbo (Bio-Rad®). We used the following antibodies: anti-CD73 (ab 133582 Abcam), anti-Vinculin (ab 129002 Abcam). We used chemiluminescence as a detection method with the Clarity Western ECL Substrate (Bio-Rad®) reagent.

### Statistical analysis

Statistical analysis and graphics were performed using GraphPad Prism® software (GraphPad Software®, Inc., La Jolla, CA, USA). For comparison between two groups, an unpaired Student’s t-test was performed. For comparison among three or more conditions, non-parametric one-way ANOVA (Mann-Whitney test) followed by Dunnett post-test was performed. A p-value of less than 0.05 was considered significant. For *in vitro* analysis, all statistical tests were based on data from at least n=3 independent experiments (experiments using independent cell batches).

## Results

### miRs associated with vascular calcification and ossification in human atherosclerotic lesions

To identify miRs associated with vascular calcification in human plaques, we performed dedicated miR microfluidic arrays that include 753 miRs on 60 arterial samples from ECLAGEN biocollection. 20 atherosclerotic carotid arteries (10 calcified, 10 none calcified), 20 atherosclerotic femoral arteries (10 ossified, 10 none ossified), 10 healthy carotid arteries, and 10 healthy femoral arteries. We identified calcification and ossification based on the presence of histological calcified structures (clear center, sheet-like, nodules, osteoid metaplasia) as previously described^11^.

To select our miR of interest associated with vascular calcification, we first compared calcified versus non-calcified carotid plaques and showed significant differences in expression for miRs 515-3p, 518e, 383, 136, 497, 548L, and 183 (Fig.1-A). We also identified miRs 548c, 1278, 640, 155, 127-5p, 554, 183, and 573 when comparing femoral plaques with osteoid metaplasia (bone-like structures) versus no osteoid metaplasia (Fig.1- B). Among these targets, we further selected the most regulated miRs and miRs differentially expressed in femoral arteries compared to carotid arteries (Fig.1-C), as we previously showed that femoral arteries were more prone to calcification compared to other arterial locations^21^. Our final selection of miR candidates included miR127, 136, 155, 183, 383, 497, 515, 518, 548, 554, 640, and 1278.

**Figure 1:**
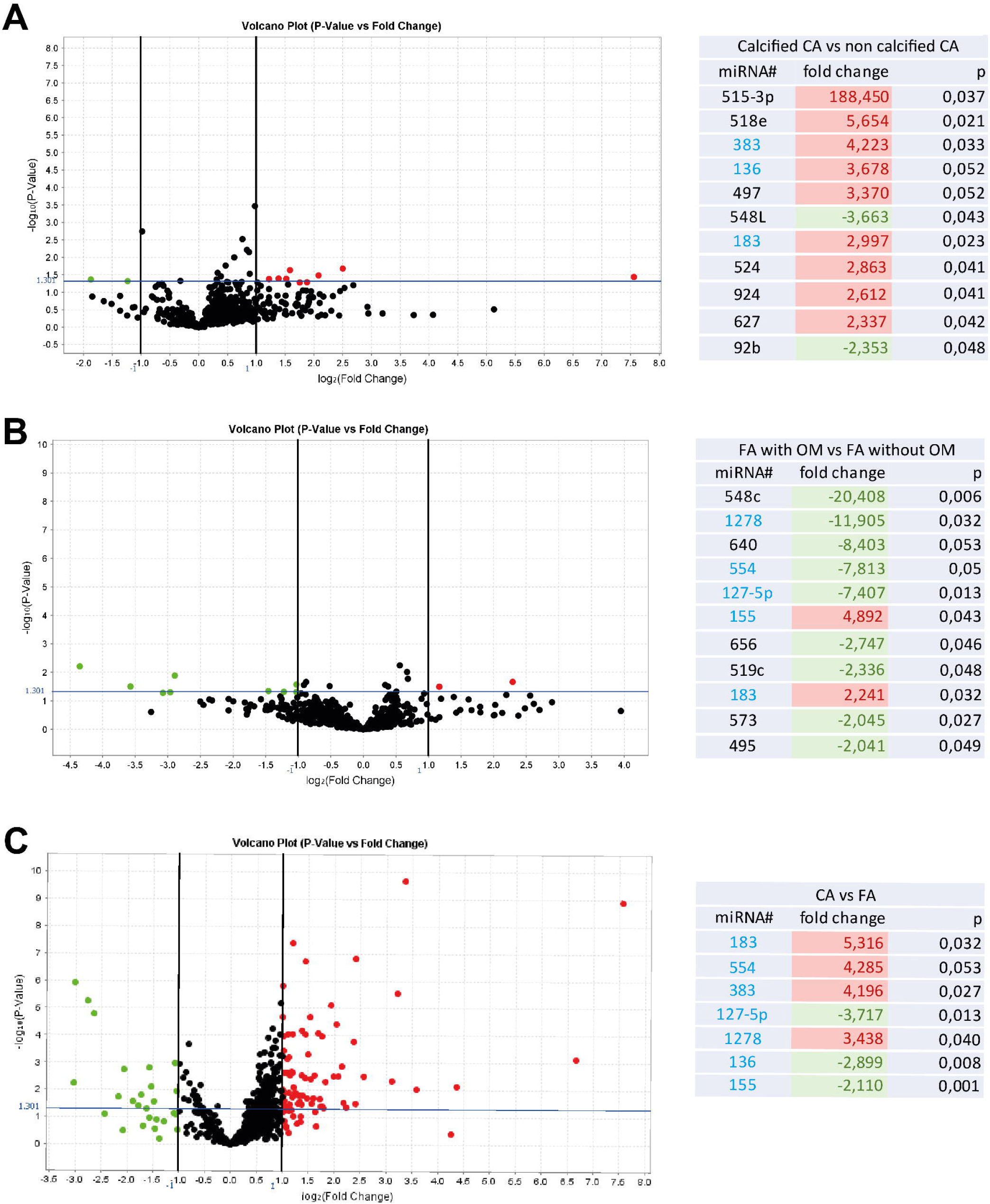
Volcano plot graphs and tables representing fold changes of miRs differentially expressed between calcified vs non-calcified pathological carotid arteries (n= 20 patients) (A), between ossified vs non-ossified pathological femoral arteries (n= 20 patients) (B), and between healthy carotid and femoral arteries (n = 10 patients)(C).

In silico analysis of the biological functions associated with the combined target genes of the 12 miR candidates showed a predominant role in smooth muscle cell lineage differentiation, including Wnt signaling and bone resorption, confirming their putative role in VSMC transition to an osteogenic phenotype. (Fig.2)

**Figure 2:**
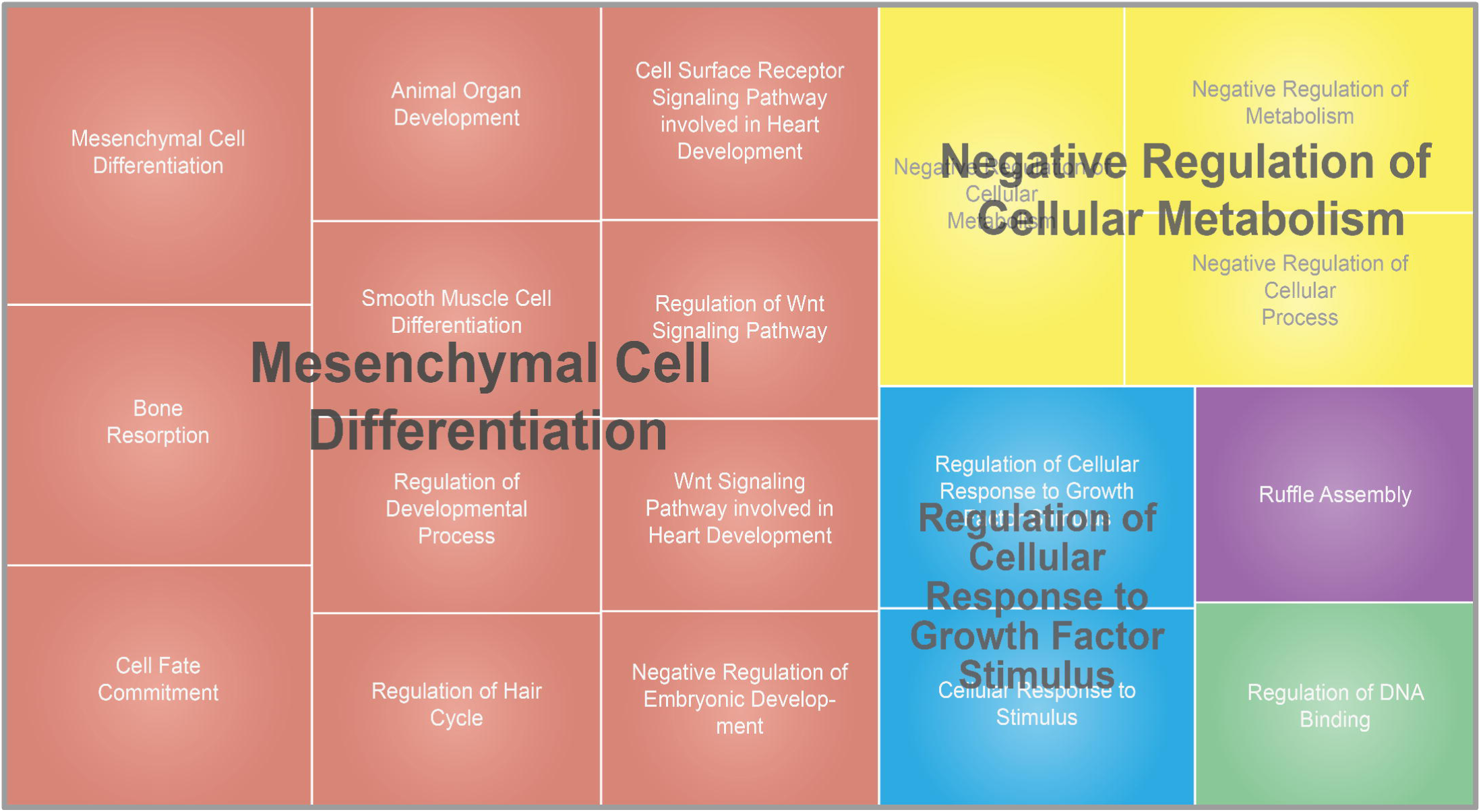
REVIGO TreeMap view of GO terms (biological functions) associated with the target genes of the 12 miR candidates. Each rectangle is a single cluster representative. The representatives are joined into ‘superclusters’ of related terms that are visualized with distinct colors (indicated by centralized black text). The size of the rectangles reflects the number of genes associated with the given biological functions.

### Vascular smooth muscle cell mineralization

#### miR expression and regulation during VSMC mineralization

VSMCs are important drivers of VC, as their phenotypic plasticity can lead to osteogenic properties. To identify miRs involved in this process, we investigated their expression in primary VSMC and their regulation during cell mineralization induced either by inorganic phosphate or by a cocktail used to induce osteoblastic differentiation from mesenchymal progenitors (beta-glycerol phosphate, dexamethasone, and ascorbic acid). We first validated the mineralization of primary human VSMC with both protocols and compared their phenotypic modulation in response to these protocols.

For inorganic phosphate-enriched medium, cell mineralization after 7 days was associated with an overexpression of Alkaline Phosphatase in early stage (2.766-fold at day 1), Osterix (2.661-fold at day 2), CollagenI-α1 (1.511-fold at day 2), Sox9 (2.251-fold at day 5), Osteocalcin (2.990 fold at day 3) and Bone Sialoprotein 2 (4.724-fold at day 1). Conversely, decrease in contractility associated genes expression of α-SMA (0.313 at day 7), Calponin-1 (0.453 at day 7), Tropomyosin-1 (0.692 at day 2), and Transgelin (0.420 at day 3) further illustrated the transition of VSMC into an osteogenic phenotype (Fig.3-A).

**Figure 3:**
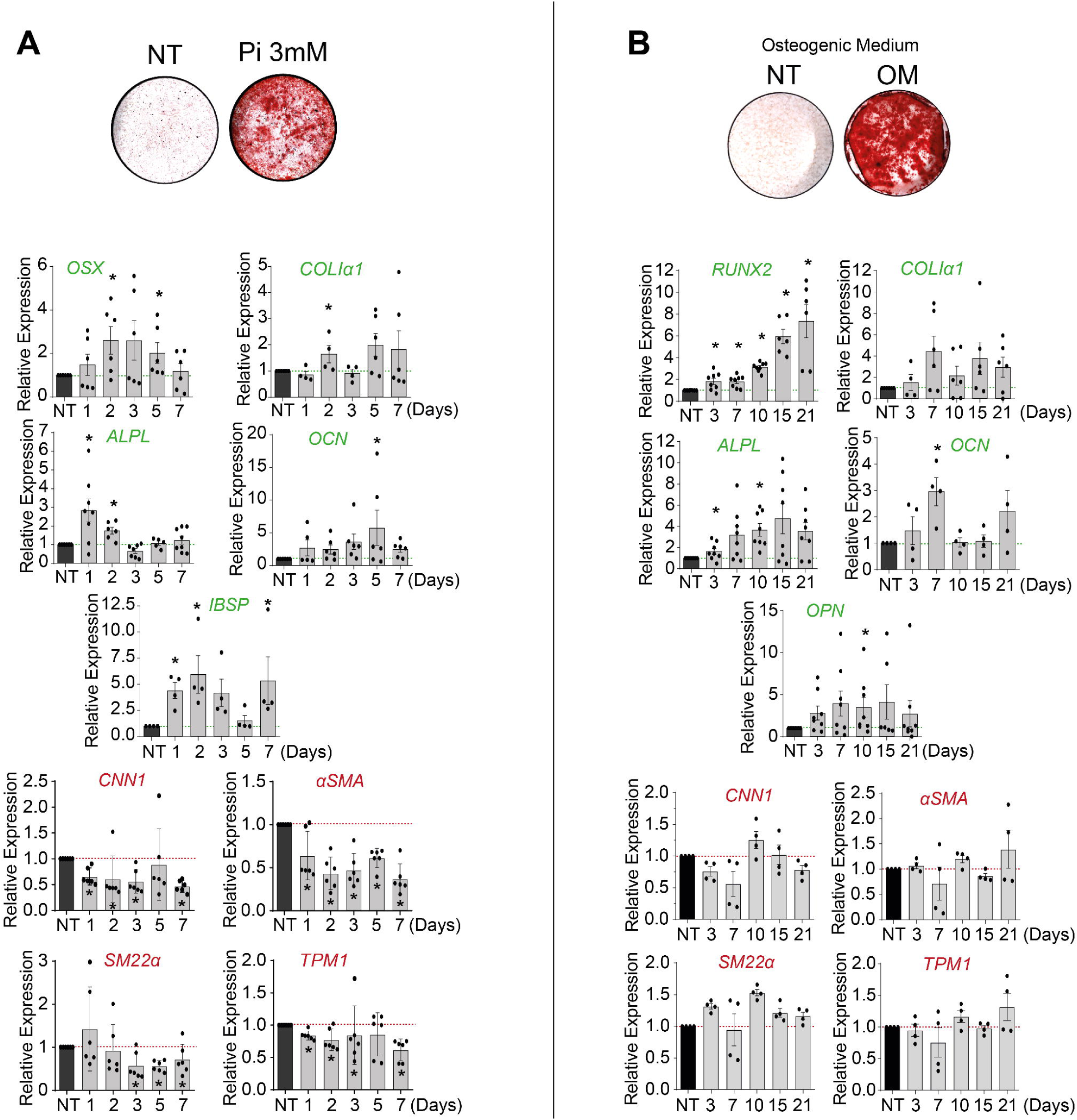
A. VSMCs mineralization stained in red with red alizarine after 10 days of culture in Inorganic phosphate enriched medium (3mM Pi). Representative image of wells after red alizarine staining. Relative RNA expression levels of osteoblastic genes (in green) and contractile genes (in red) during VSMC mineralization. B. VSMCs mineralization after 21 days in culture in Osteoblastic Medium. Representative image of wells after red alizarine staining. Relative RNA expression levels of osteoblastic genes (in green) and contractile genes (in red) during VSMC mineralization. Bars represent mean ± SEM (*p<0.05).

For osteoblastic medium (OB), mineralization obtained after 21 days was also correlated with overexpression of Alkaline Phosphatase (3.183-fold at day 10), Osteopontin (1.873-fold at day 10), and Osteocalcin (3.064-fold at day 7). We also found a tendency of an overexpression of CollagenI-α1. In contrast to inorganic phosphate, this osteogenic medium increases Runx2 expression throughout the treatment (8.651-fold at day 21) (Fig.3-B).

Analysis of the 12 miRs of interest expressed in both pro-mineralizing conditions allowed us to exclude eight miRs that were either not expressed in VSMCs or not regulated during VSMC mineralization (miRs 383, 497, 515, 518, 548, 554, 640, 1278). During Pi-driven mineralization, we observed some regulations for 3 miR candidates (miR136, 155, and 183). An increased expression of miR136 (1.828 fold at day 2), miR155 (2.381 fold at day 7), and miR183 (4.53 fold at day 5) was apparent (Fig.4-A). OB medium also regulated transiently 4 miRs of interest. We observed an up-regulation of miR136 (2.346-fold at day 7), miR155 (5.242-fold at day 7), and miR183 (3.45-fold at day 7). Conversely, we observed a down- regulation of miR127 (0.357 at day 21) (Fig.4-B).

**Figure 4:**
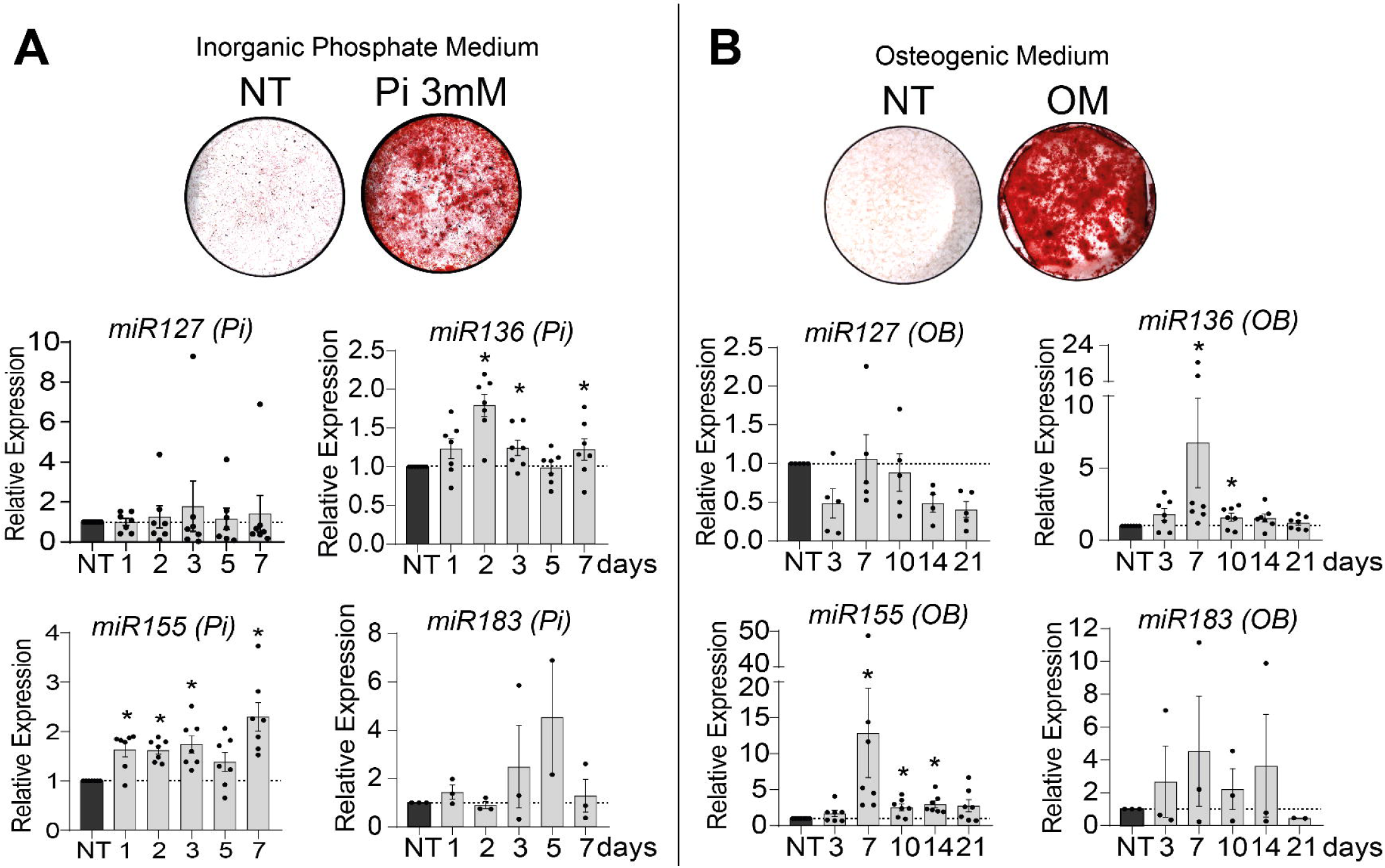
miR expression levels during VSMCs mineralization induced by inorganic phosphate (3mM) (A) and Osteoblastic Medium (B) for each miR candidate: 127, 136, 155, and 183. Bars represent mean ± SEM (*p<0.05).

#### Functional analysis of the effect of each targeted miR

To assess their functional importance in VSMC mineralization, we overexpressed miRs by lipofection for the duration of the process. Using this approach, miR overexpression was maintained for 7 days after transfection (Fig.5-A).

**Figure 5:**
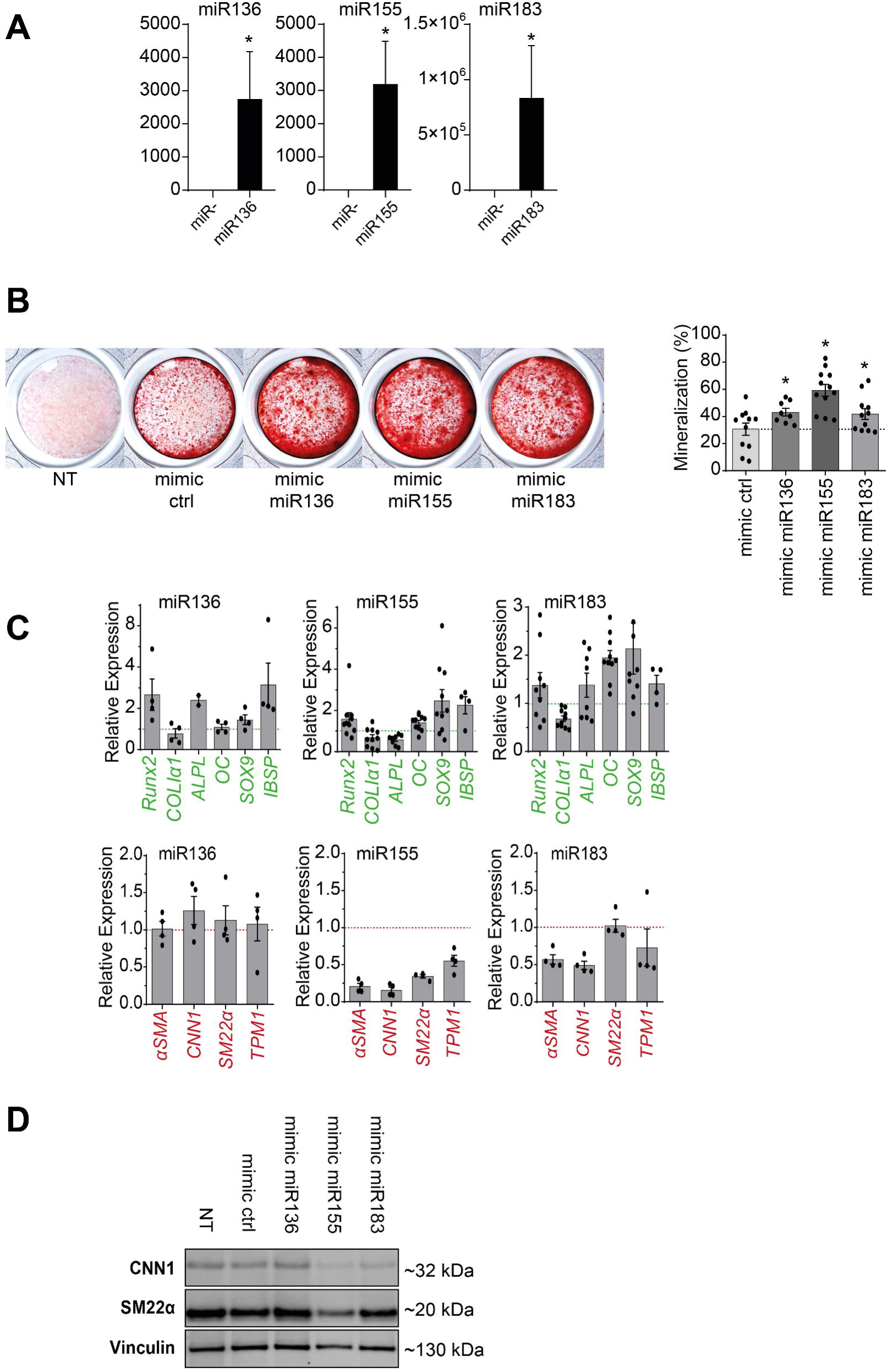
Functional impact of miR candidates on vascular smooth muscle cell mineralization. A: miRs 136, 155, and 183 overexpression in VSMCs 7 days after transfection of individual miR mimics. B: Representative images of alizarin red staining for mineralization (in red) of VSMC 7 days after inducing miR overexpression (mimics negative control, miR136, miR155, and miR183) in Pi-enriched medium (3mM Pi), and quantification of the mineralized area percentage (stained area/total area) between conditions. C: Transcriptional regulation of osteoblastic (in green) and contractile-associated genes (in red) 7 days after inducing miR overexpression compared to a negative ctrl mimic. Bars represent mean ± SEM (*p<0.05). D: Representative western blotting for contractile-associated proteins 7 days after miR mimics transfection.

Analysis of miRs overexpression on VSMC mineralization (Fig.5-B) showed a significant increase of the mineralization with the miR155 compared to control (59.6 vs 37.8%, p=0.0007). We observed a minor increase in mineralization after transfection of mimics miR136 and miR183. Transfection of mimic miR127 did not result in any increase in cell mineralization over mimic control.

Overexpression of miR155 in phosphate-enriched mineralization medium led to a significant increase in the expression of several osteoblastic genes Runx-2 (1.353-fold, p=0.014), Sox-9 (1.914-fold, p=0.014), Osteopontin (1.361-fold, p=0.008), Osteocalcin (1.532-fold, p=0.0006) and Bone Sialoprotein-2 (2.551-fold, p=0.027). We also observed a significant decrease in the expression of the CML-specific markers α-SMA (0.195, p=0.029), calponin (0.154, p=0.029), transgelin (0.359, p=0.029), and tropomyosin-1 (0.561, p=0.029) (Fig. 5-C), which was confirmed at the protein level by Western blotting (Fig. 5-D).

The over-expression of miR136 also resulted in a significant increase in the expression of osteoblastic genes Runx-2 (2.181-fold, p=0.001), Alkaline Phosphatase (2.391-fold, p=0.022), and Bone Sialoprotein-2 (2.159-fold, p=0.029) but we did not observe any significant decrease in contractile VSMC genes expression (Fig.5-C).

The over-expression of miR183 also resulted in a minor but significant increase in the expression of osteoblast genes osteocalcin (1.876-fold, p=0.0001), and Sox-9 (1.608-fold, p=0.001). Conversely, there was a significant decrease in Collagen-1 (0.647, p=0.0001) and VSMC markers α-SMA (0.519, p=0.029) and Calponin (0.455, p=0.029) (Fig.5-C and Fig.5- D).

Among these miR candidates, miR155 appeared to play the most effective role in VSMC mineralization and osteogenic trans-differentiation.

### Associated miR155 targeted genes identification

To uncover the genes targeted by these miRs implicated in vascular calcification in human lesions, we listed all the target genes from those miRs (miRWalk 2.0, miRDB, miRbase Databases) and crossed our transcriptomic analysis from the biocollection to identify genes related to tissue mineralization and bone metabolism with a regulation consistent with miR expression in calcified lesions (Fig.6 and Fig.7).

**Figure 6:**
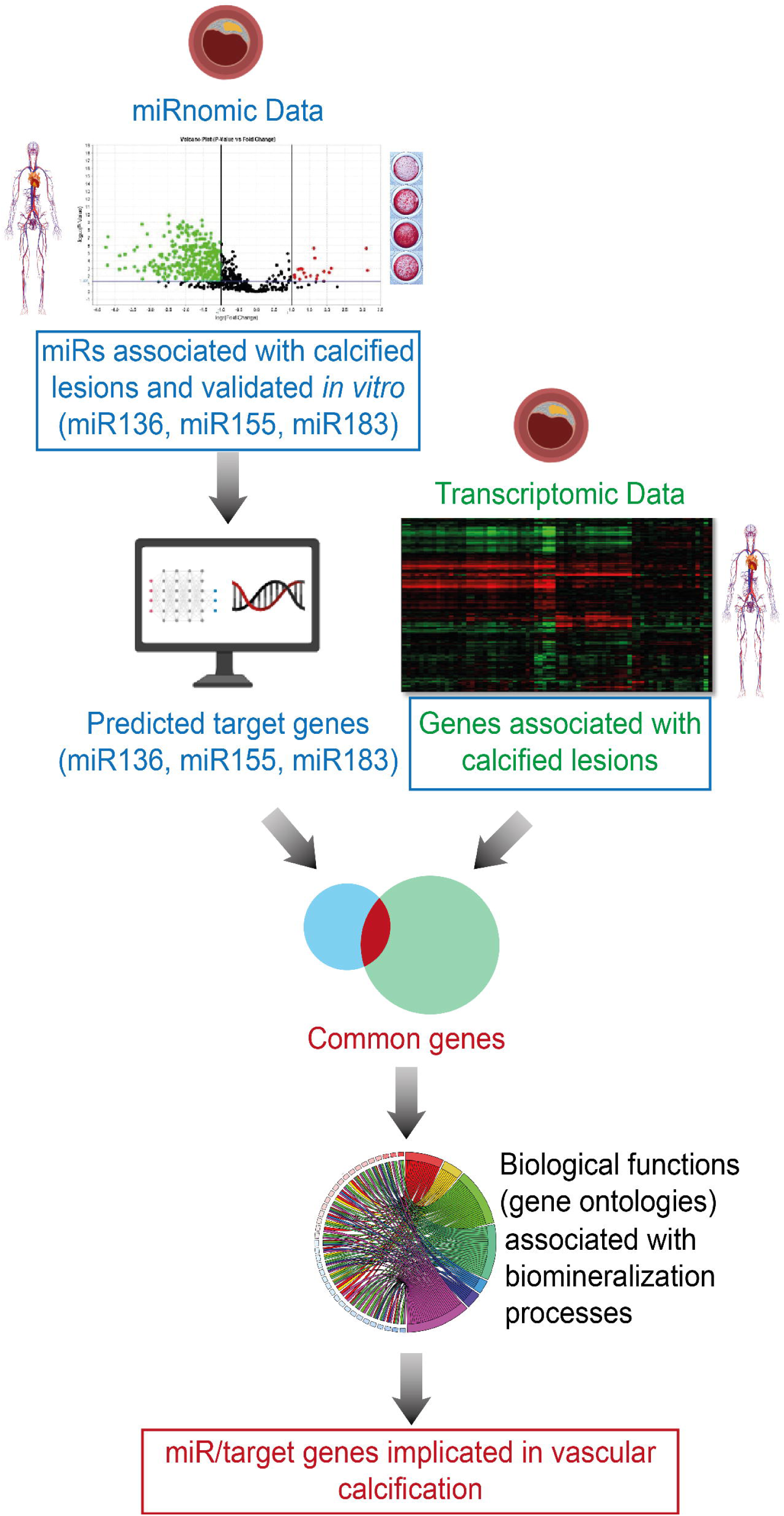
Diagram illustrating our strategy consisting in crossing our miRnomics and our transcriptomic data for the identification of miRs/target genes associated with vascular calcification (VC) in ECLAGEN human biocollection samples. In silico analysis predicted the putative target genes of our miRs of interest (miR Walk, miRDB, miRbase Databases). Among them, we selected those with a consistent regulation to their respective miR in calcified lesions compared to none-calcified lesions. Finally, a gene ontology analysis (biological function analysis) of each of these target genes allowed the selection of our miR putative target-genes with a known involvement in biomineralization process.

**Figure 7:**
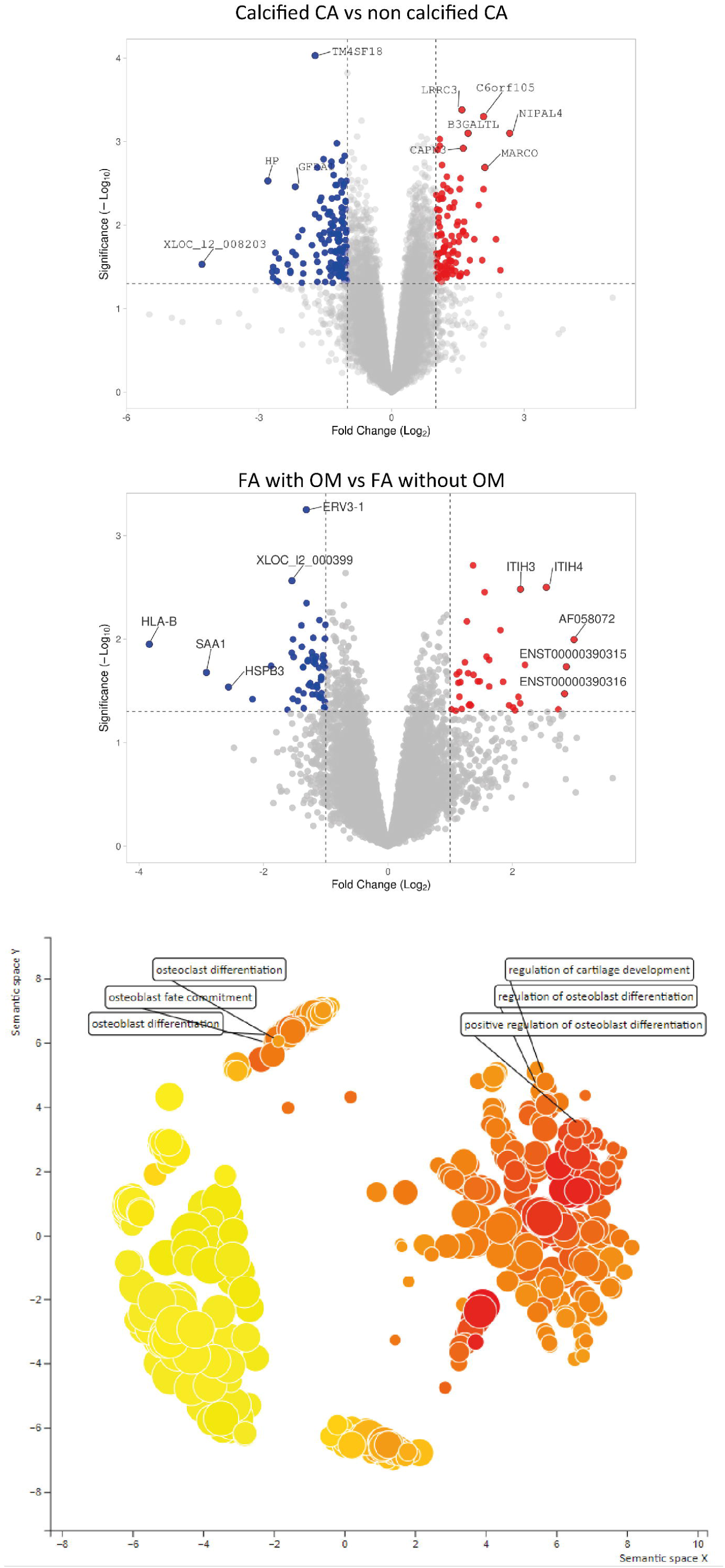
Volcano plot graphs representing fold changes of genes differentially expressed between calcified vs non-calcified pathological carotid arteries (n=20) (A), and between ossified vs non-ossified pathological femoral arteries (n= 20) (B). REVIGO Scatterplot of their gene ontology annotations (biological functions) highlighting their contributing roles in biomineralization process (C).

Putative target genes of our miR of interest regulated in calcified and ossified lesions were illustrated in Fig.7-A and Fig.7-B. A gene ontology analysis of their functional annotations highlighted their contributing roles in biomeralization (Fig.7-C). This strategy aiming to highlight miR/target genes duplexes associated with VC led to the identification of 4 genes: Smad3, Fli1, CD73, and Wnt5a. Consistently with our projection, Smad3 can play an inhibitory role in phosphate-induced vascular smooth muscle cell calcification^28^, and CD73 can down-regulate TNAP activity, a major enzyme that regulates the extra-cellular concentration of Pyrophosphate (a potent inhibitor of VC)^29^.

To confirm the link between miR155 and these genes that may control its pro-mineralizing properties, we examined the transcriptional and protein expression regulation of CD73 and Smad3 on VSMC after transfection of miR155 (Fig. 8). The transcriptional study showed a significant decrease in CD73 and Smad3 gene expression induced by miR155 compared to control (0.630, p<0.0001 and 0.770, p<0.0001, respectively). This was further confirmed at the protein level for CD73 (0.754, p=0.0002).

**Figure 8:**
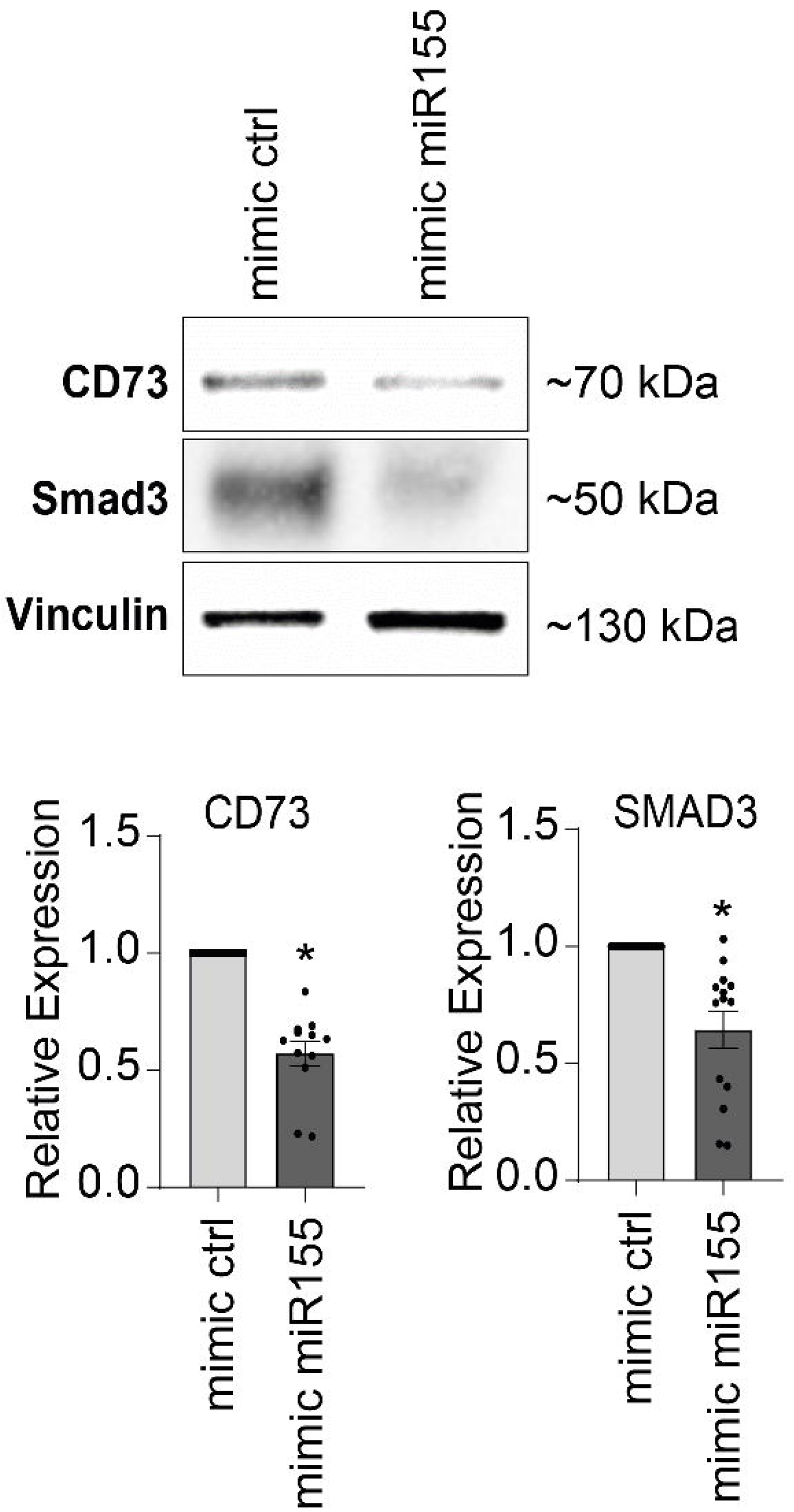
Smad3 and CD73 protein and RNA levels 2 days following overexpression of indicated miR mimics. Bars represent mean ± SEM (*p<0.05)

## Discussion

In the present study, we identified miRs that are associated with vascular calcification in human atherosclerotic lesions and that participate in vascular smooth muscle cell mineralization *in vitro*. Other studies have investigated the expression profile of miRs in human atherosclerotic plaques and VC. Raitoharju et al., based on the Tampere Vascular Study, compared miRs expression in atherosclerotic plaques from carotid, femoral, and aortic arteries. The atherosclerotic plaques were compared with samples from the territory of the internal mammary artery, which is known for its low susceptibility to atherosclerosis. This work showed a significant up-regulation of the expression of miR21, -34a, -146a, -146b-5p, and 210 (4.61, 2.55, 2.87, 2.82, and 3.92-fold changes, respectively)^30^. Some of them have been associated in vitro with VSMC mineralization^31^. In parallel, recent studies have identified miRs that regulate vascular calcification in different models, both *in vivo* and *in vitro*. For example, miR16-5p, miR17-5p, miR20a-5p, miR106b-5p, and miR204/miR-211 have been described as protective against medial calcification, and are depleted in extracellular vesicles in CKD patients^32,33^. Similarly, miR126-3p, miR33a-5p, and miR223-3p have been shown to reduce calcification in aortic valves in murine experimental models^34–36^. In addition to these studies, emerging data highlight the predictive value of miRs in CVD. For VC, serum miR-125b levels associated with VC severity could be used to reflect uremia- associated calcification in CKD patients^37^.

Our study focuses on intimal calcification, a very common and heterogeneous form of VC. We employed a non-biased strategy to identify miRs associated with VC in carotid and femoral arteries. By considering both carotid calcifications and femoral ossifications, our initial objective was to identify miRs that may be involved in these two major types of mineralization and territories. We assumed that common factors might still emerge (like miR183). However, only few common miRs were identified. This led to the identification of miRs that could play a significant role in vascular calcification, but maybe at different stages of their development. This strategy allowed us to identify a dozen miRs enriched or depleted in calcified or ossified lesions, so potentially important in vascular calcification, but maybe at different stages of their development. After a validation step *in vitro*, we showed that most of these miRs were not expressed in primary human VSMC, indicating that they may also act indirectly in VC, potentially by regulating endothelial, immune, and/or fibroblast cell biology that could affect VC. They could also be expressed by more differentiated local osteoblastic cells that we don’t replicate in vitro with mineralizing vascular SMC. The cellular origin of these miRs and their functional relevance in VC will be investigated in future studies.

This strategy also aimed to identify miRs involved in various forms of VC by examining both the carotid and femoral arteries, which can host different types of VC^11^. We also assessed miRs expression during VSMC mineralization using two protocols widely used in the literature. Inorganic phosphate and osteogenic cocktail can both trigger VSMC mineralization, with specific kinetics and molecular pathways^38^. One is commonly used as a model for VSMC mineralization as seen in patients with chronic kidney disease, associated with hyperphosphatemia and medial calcification, and the latter is employed to mimic osteoblast differentiation from mesenchymal progenitors. In this study, we selected miRs that were commonly regulated during mineralization with each protocol, suggesting that they could act as a shared molecular player in these processes.

To date, there are still no dedicated pharmacological therapeutic strategies targeting VC. In our study, miR155 had the most pronounced pro-mineralizing property compared to miR136 and miR183. This confirms recent studies reporting the involvement of miR155 in VC *in/ex vivo*. Using murine aorta rings, miR155 deficiency attenuated calcification, and miR155^-/-^ mice showed reduced VC induced by vitamin D3^39^. Moreover, Zhao et al. found an important regulatory role for the RCN2/STAT3/miR155 loop in hyperphosphatemia-induced VC, particularly in patients with chronic renal failure^40^. Finally, the study by Fakhry et al. also reported the overexpression of miR155 in phosphate-induced rat aortic mineralization^41^.

In addition to VC, miR155 plays a critical role in the regulation of the inflammatory response, and thus in atherosclerosis^42^. Elevated miR155 levels have been associated with inflammatory macrophages and atherosclerotic lesions, but its effects may differ in early and advanced atherosclerosis depending on plaque content^43,44^. Nonetheless, inhibition of miR155 in immune cells appears to be a promising strategy to limit the development of atherosclerosis^45^, certain cancers^46^, and inflammatory diseases such as rheumatoid arthritis^47^. Our work further extends this list, by highlighting the potential benefit of miR155 inhibition in limiting VC development in peripheral atherosclerotic arteries, but additional experiments with miR155 targeting already diseased and calcified arteries need to assess its potential clinical value. We will explore the impact of these miRs in early and late-stage calcification, and overall, test whether miR targeting can reduce or limit already developed mineralization, directly or/and indirectly, through a local or systemic approach.

## Acknowledgments and Funding

We are most grateful to the GenoBIRD Core Facility for its technical support. We thank Carine Montagne, Flavien Gautron, and Manon Pondjikli for the management of biocollections, which was funded by the Allocation Nationale de Recherche (ANR) for physiopathology and by an inter-regional Programme Hospitalier de Recherche Clinique (PHRC).

This work was funded by the Fondation de l’Avenir (Paris, France), the University Hospital of Nantes (Nantes, France), the Fédération Française de Cardiologie (Paris, France), the Société Francaise de Cardiologie (Paris, France), and the Societé de chirurgie vasculaire et endovasculaire (Paris, France).

## Contributions

Tom Le Corvec: Investigation, Formal analysis, Writing - Original Draft Visualization; Mathilde Burgaud: Investigation, Validation; Marja Steenman: Data Curation; Robel A Tesfaye: Data Curation; Yann Goueffic: Conceptualization, Funding acquisition; Blandine Maurel-Desanlis: Writing - Review & Editing Conceptualization, Writing- Reviewing and Editing, Supervision, Funding acquisition; Thibaut Quillard: Writing - Review & Editing Conceptualization, Writing- Reviewing and Editing, Visualization, Supervision, Project administration, Funding acquisition.

